# A folded viral noncoding RNA blocks host cell exoribonucleases through programmed remodeling of RNA structure

**DOI:** 10.1101/262683

**Authors:** Anna-Lena Steckelberg, Benjamin M. Akiyama, David A. Costantino, Tim L. Sit, Jay C. Nix, Jeffrey S. Kieft

## Abstract

Folded RNA elements that block processive 5′→3′ cellular exoribonucleases (xrRNAs) to produce biologically active viral non-coding RNAs were discovered in flaviviruses, potentially revealing a new mode of RNA maturation. However, it was unknown if this RNA structure-dependent mechanism exists elsewhere and if so, whether a singular RNA fold is required. Here, we demonstrate the existence of authentic RNA structure-dependent xrRNAs in dianthoviruses, plant-infecting viruses unrelated to animal-infecting flaviviruses. These novel xrRNAs have no sequence similarity to known xrRNAs, thus we used a combination of biochemistry and virology to characterize their sequence requirements and mechanism of stopping exoribonucleases. By solving the structure of a dianthovirus xrRNAs by x-ray crystallography, we reveal a complex fold that is very different from the flavivirus xrRNAs. However, both versions of xrRNAs contain a unique topological feature that is created by a different set of intramolecular contacts; this may be a defining structural feature of xrRNAs. Remarkably, the dianthovirus xrRNA can use ‘co-degradational remodeling,’ exploiting the exoribonuclease’s degradation-linked helicase activity to help form their resistant structure; such a mechanism has not previously been reported. Convergent evolution has created RNA structure-dependent exoribonuclease resistance in different contexts, which establishes it as a general RNA maturation mechanism and defines xrRNAs as an authentic functional class of RNAs.

## Introduction

During eukaryotic cellular RNA decay, 5′→3′ hydrolysis by an XRN protein is important for degrading decapped messenger RNA (mRNA) and other 5′-monophosphorylated RNAs including fragments of endonucleotyic cleavage, ribosomal RNAs (rRNA), and transfer RNA. Xrn1 is the dominant cytoplasmic 5′→3′ exoribonuclease in most eukaryotic cells (1, 2), where its processive translocation-coupled unwinding of RNA helices allows it to efficiently hydrolyze structured RNA substrates without releasing partially degraded intermediates (3, 4). Xrn1 plays a central role during constitutive mRNA turnover and RNA quality control, and it has been implicated in degrading viral RNAs as part of the cell’s antiviral response (5). Despite the efficiency of Xrn1, some viruses have evolved RNA sequences that robustly block the enzyme’s progression. This ability is conferred by their folded three-dimensional structures without the help of accessory proteins, thus we refer to them as Xrn1-resistant RNAs (xrRNAs) (6, 7). xrRNAs were originally identified in the positive-sense RNA genomes of mosquito-borne flaviviruses (e.g. Dengue virus, Zika virus, West Nile virus) where they protect the viral genome’s 3′ untranslated region (UTR) from degradation, generating biologically active non-coding RNAs involved in cytopathic outcomes and pathogenicity during infection (8–17). Thus, these mosquito-borne flaviviruses usurp Xrn1’s powerful degradation activity as part of an elegant RNA maturation pathway that relies on a compact and unusual RNA fold. Specifically, these xrRNAs form an interwoven pseudoknot conformation that is centered on a conserved three-helix junction, creating a protective ring-like structure that wraps around the 5′ end of the xrRNA (7, 18). This unique topology acts as a mechanical block to Xrn1 when the ring-like structure braces against the surface of the enzyme (7, 19, 20). To date, this topology and resultant mechanism has only been observed in the flaviviruses.

Recruiting Xrn1 to a larger precursor RNA, then blocking the enzyme using a compact structured RNA element to generate a biologically active smaller RNA could represent a useful general mechanism for RNA maturation. Consistent with this, xrRNAs have been identified in a broad range of flaviviruses including those that are tick-borne, those specific to insects, and those with no known arthropod vector (20). Based on secondary structural patterns, flaviviral xrRNAs can be grouped into two classes: all xrRNAs from mosquito-borne flaviviruses identified to date belong to class 1 xrRNAs, others harbor class 2 xrRNAs (20). The three-dimensional structure of a class 2 flavivirus RNA has not been solved, thus for simplicity we will use the term ‘flavivirus xrRNA’ here to refer to the well-characterized class 1 flavivirus xrRNAs, unless otherwise specified. In addition, RNA sequences that appear to resist 5′→3′ exoribonuclease degradation have been identified in a few other virus families (21–24), However, nothing is known about the molecular processes or structures of putative xrRNAs outside the flavivirus family. We do not know if other candidate exoribonuclease–resistant RNAs operate with the help of protein factors or if they are stably structured elements sufficient to block processive degradation. If RNA structure plays a role in diverse putative xrRNAs, we do not know whether there is a universal structural feature that confers Xrn1 resistance or many structures that can block Xrn1. The degree to which different xrRNAs operate by creating a mechanical block for Xrn1 versus using some other means to stop the enzyme remains unexplored. Indeed, if viruses outside the flavivirus family use diverse sequences and structures to block exoribonucleases in a programmed way, this would indicate that this mechanism is a general pathway and that xrRNAs are a true functional class of RNAs.

To address these questions, we investigated an unexplored potential xrRNA sequence from the 3′UTR of plant-infecting dianthoviruses (25, 26) (Fig. S1*A*). Using a combination of virology and biochemistry, we show that the dianthovirus 3′UTRs contain a *bona fide* xrRNA sequence that inhibits 5′→3′ exoribonucleolytic decay *in cis* without the help of protein factors. X-ray crystallography and single-molecule FRET experiments reveal a unique RNA fold that operates to block 5′→3′ exoribonucleolytic degradation through dynamic changes in RNA structure that we term ‘co-degradational’ RNA remodeling. This work reveals RNA-dependent exoribonuclease resistance beyond the realm of flaviviruses, conferred by diverse structured RNA elements that nonetheless share a common underlying topology. Thus, convergent evolution has given rise to this mechanism at least twice, indicating it to be a generally useful pathway that may be widespread across biology.

## Results

### The dianthovirus 3′UTR contains a *bona fide* xrRNA

We verified that dianthovirus infection results in accumulation of a subgenomic RNA by infecting *Nicotiana benthamiana* with Red clover necrotic mosaic virus (RCNMV) (27). Northern blot analysis of RNA from infected leaves with probes targeting the viral 3′UTR revealed the presence of a discrete RNA species consistent with the previously reported SR1f RNA (Fig. 1*A*), which has been shown to be the result of 5′→3′ exoribonuclease resistance. To determine if RNAs with sequences from the dianthovirus 3′UTR are sufficient to block 5′→3′ exoribonucleolytic decay without *trans-acting* proteins, we challenged *in vitro*-transcribed RNA from RCNMV with recombinant Xrn1 (Fig. S1*B*). A 130 nucleotide-long RNA from the RCNMV 3′UTR blocked exoribonucleolytic decay with no protein cofactors, revealing that the RCNMV contains an authentic xrRNA (Fig. 1B). We mapped the Xrn1 halt site using reverse transcription and capillary electrophoresis (Fig. S1*D*-*F*), observing that the enzyme halts at a specific point. We then determined the 3′ terminus of the resistant sequence using test RNAs truncated at their 3′ end, identifying a 43-nt RNA element as the minimal RCNMV xrRNA (Fig. 1*B*). Thus, the RCNMV 3’UTR has an authentic xrRNA contained in a discrete RNA sequence that can operate outside of its natural context.

**Fig. 1.**
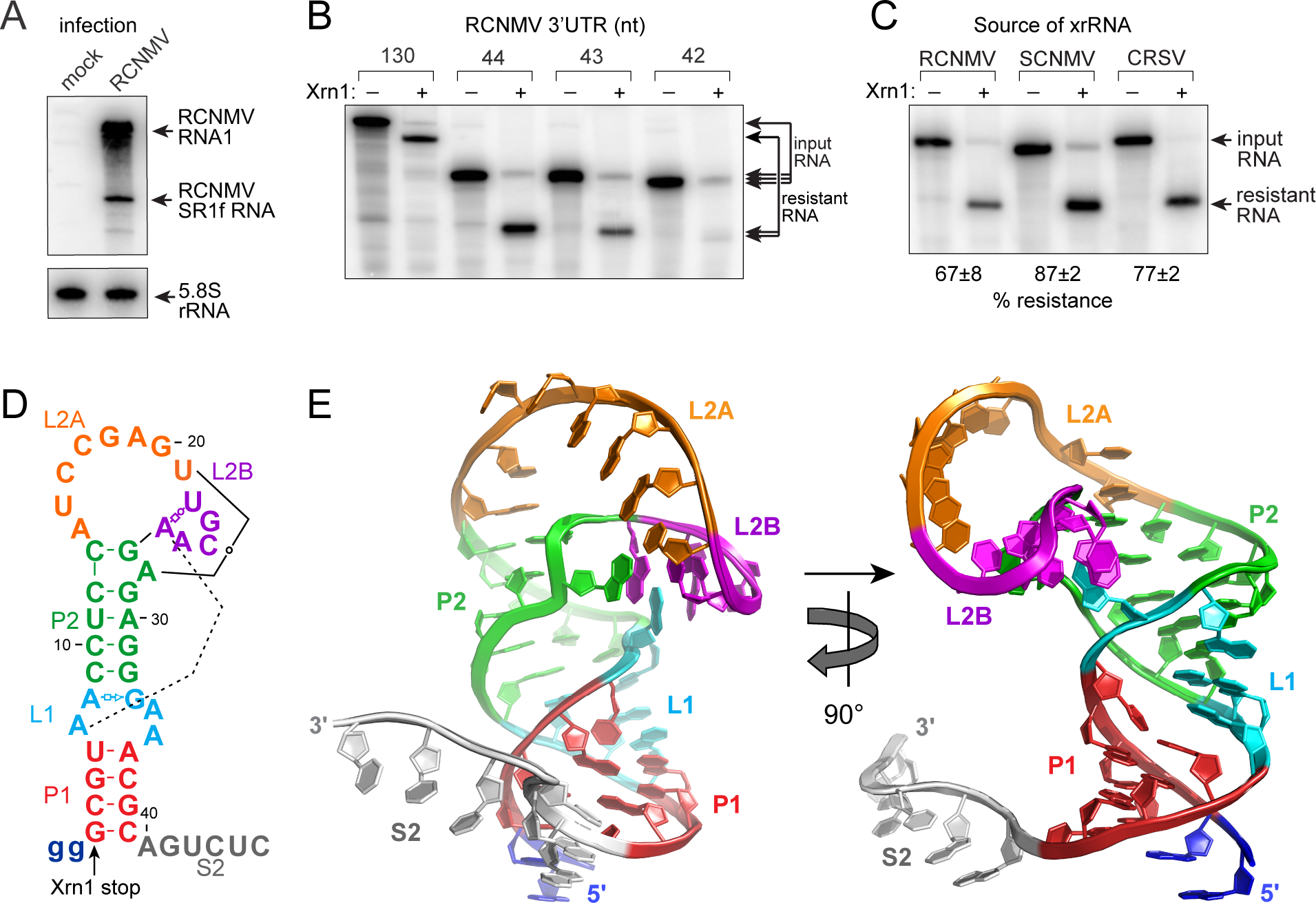
Structure of an authentic xrRNA in dianthovirus 3′UTRs. (*A*) Northern blot of total RNA from mock- or RCNMV-infected *N. benthamiana*. Probes are against viral genomic 3′UTR and 5.8S rRNA. Full-length RNA1 and exoribonuclease-resistant degradation product (SR1f) are indicated. (*B*) *In vitro* Xrn1 degradation assay of ^32^P-3′-end-labeled RCNMV 3′UTR sequences. (*C*) *In vitro* Xrn1 degradation assay on minimal xrRNAs from RCNMV, SCNMV, and CRSV. Average % resistance from 3 individual experiments (-/+ SD). (*D*) Secondary structure of the crystallized SCNMV RNA. Lowercase letters represent sequences altered to facilitate transcription. Non-Watson-Crick base pairs in Leontis-Westhof annotation (47). Xrn1 stop site: labeled arrow. (*E*) Ribbon representation of the SCNMV xrRNA structure. Colors match (*D*).

The xrRNA sequence identified in RCNMV is highly conserved in all three known members of the dianthovirus family (Fig. S1*C*) (RCNMV; Sweet clover necrotic mosaic virus, SCNMV; Carnation ringspot virus, CRSV) and accordingly, all viral 3′UTR sequences confer Xrn1 resistance *in vitro* (Fig. 1*C*). As the mosquito-borne flavivirus xrRNAs are able to block diverse exoribonucleases and thus act as general mechanical blocks to these enzymes (20), we tested the three dianthovirus xrRNAs for this ability. All three blocked two 5′→3′ exoribonucleases unrelated to Xrn1 (i.e. yeast decapping and exoribonuclease protein 1 (Dxo1) and bacterial 5′→3′ exoribonuclease RNase J1), indicating that they function as general roadblocks against exoribonucleolytic degradation rather than through specific interactions with the enzyme (Fig. S1*G-I*). That this matches the behavior of flavivirus xrRNAs suggests potential similarity in the molecular mechanisms of these xrRNAs. This result also justifies the use of recombinant yeast Xrn1 in our experiments as a proxy for the larger group of eukaryotic XRN proteins, which is important because dianthoviruses infect plant cells whose major cytoplasmic 5′→3′ exoribonuclease is Xrn4 (2).

### Dianthovirus xrRNAs adopt an unexpected structure

Dianthovirus xrRNAs are significantly shorter than flaviviral xrRNAs and there is no readily identified sequence similarity between the two, suggesting that a different structure confers Xrn1 resistance. To understand the structural basis of Xrn1 resistance by the dianthovirus xrRNAs, we solved the structure of the complete 43-nt SCNMV xrRNA to 2.9 Å resolution using x-ray crystallography (Table S1, Fig. S2). The SCNMV xrRNA adopts a global conformation with no immediate similarity to the mosquito-borne flaviviral xrRNA structures (7, 18). Overall, the SCNMV xrRNA structure comprises a stem-loop (SL) structure with a single-stranded 3′ tail (Fig. *2D, E*). The stem contains coaxially positioned helices P1 and P2, which are linked by stacking of bases and a noncanonical base pair in the L1 internal loop. Helix P2 is capped by L2, an atypical bipartite loop. The 5′ portion of the loop contains a run of single-stranded stacked bases (L2A) followed by a sharp turn in the backbone and a compact hairpin structure (L2B) that is embedded within the larger loop structure. This L2B element forms long-distance tertiary interactions with the L1 internal loop, causing L2 to adopt an overall ‘tilted over’ conformation. Sequence conservation between the dianthoviral xrRNAs suggests that this structure is representative of all three known members (Fig. S1*C*).

**Fig. 2.**
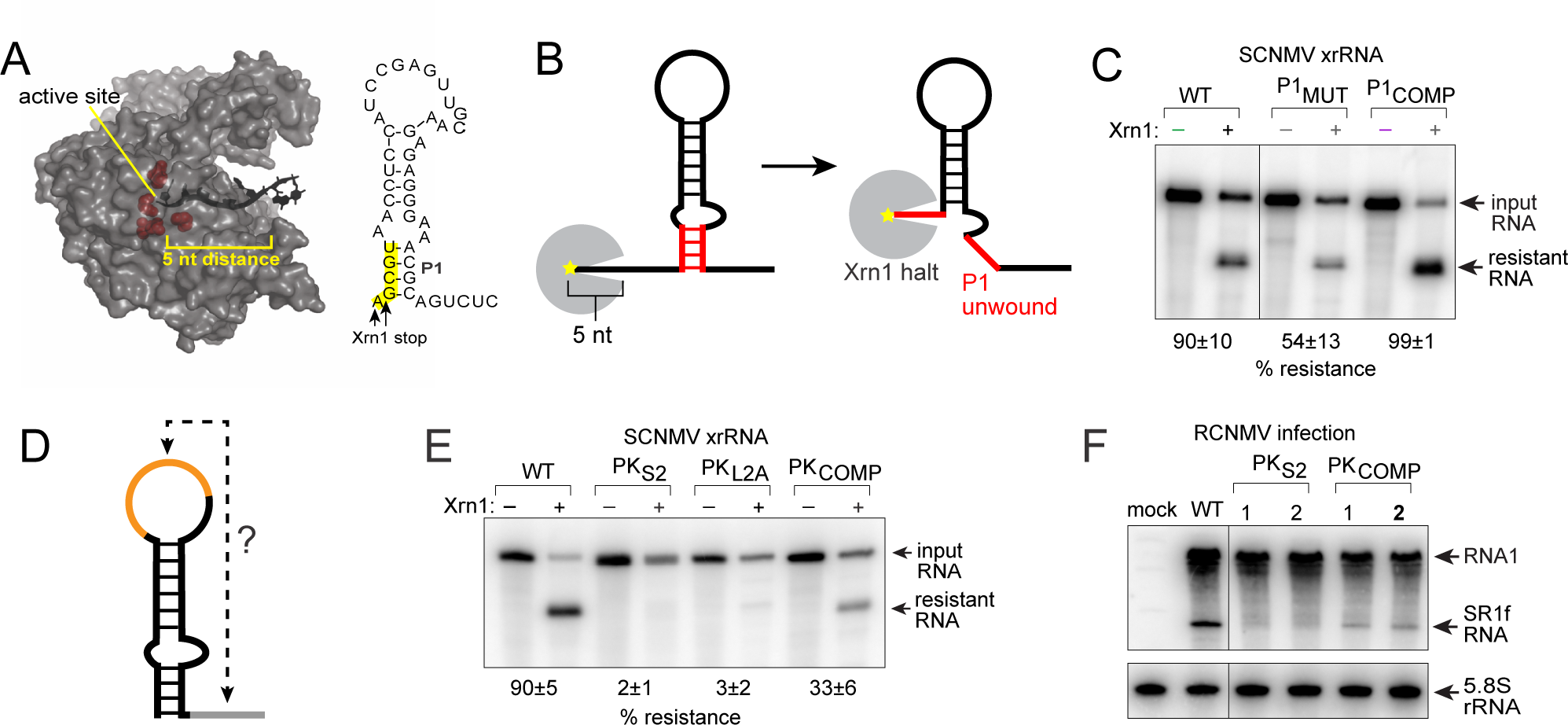
Partial unfolding of SCNMV xrRNA contributes to Xrn1 resistance. (*A*) Crystal structure of *D. melanogaster* Xrn1 (PDB: 2Y35) (3). Active site residues are depicted as red spheres and an 8-nt RNA substrate is shown in black. Five single-stranded nucleotides span the distance from the active site to the enzyme’s surface. (*B*) Scheme of Xrn1 unwinding the P1 stem (depicted in red). (*C*) *In vitro* Xrn1 degradation assay on ^32^P-3′-end-labeled WT and mutant xrRNAs. Average % resistance from 3 individual experiments (-/+ SD). (*D*) Scheme of putative pseudoknot interaction suggested by crystal contacts (see also Fig. S4). (*E*) *In vitro* Xrn1 degradation assay on ^32^P-3′-end-labeled WT and mutant xrRNAs. Average % resistance from 3 individual experiments (-/+ SD). *(F)* Northern blot of total RNA from mock- or RCNMV-infected *N. benthamiana*. Probe was against RCNMV 3′UTR or 5.8S rRNA.

### The dianthoviral xrRNA crystal structure is an RNA folding intermediate

The SCNMV xrRNA structure did not immediately suggest a mechanism for Xrn1 resistance, but functional data provided insight. The Xrn1 stop site is at the base of the P1 stem (Fig. 1*D*). This is significant because the entry channel into Xrn1’s active site only accommodates single-stranded RNA, mandating that the helicase activity of the enzyme unwinds RNA so it can enter the enzyme’s interior. The distance from the surface of the enzyme to the buried active site requires that at least 5-6 nucleotides of RNA downstream of the stop site must be single-stranded (Fig. 2*A*) (3, 4). Thus, the location of the Xrn1 stop site on the dianthovirus xrRNA mandates that P1 is unwound at the moment that Xrn1 halts (Fig. 2*B*). This raised the question of whether formation of the P1 stem is needed to block the enzyme. We tested this by measuring Xrn1 resistance of an RNA mutated to disrupt P1 base pairing (P1_MUT_) (all mutants: Fig. S3). This mutation significantly reduced but did not abolish Xrn1 resistance *in vitro*, while compensatory mutations (P1_COMP_) that restore Watson-Crick base pairing completely rescued Xrn1 resistance (Fig. 2*C*). Thus, formation of the P1 stem is not essential for Xrn1 resistance but contributes to full activity, yet P1 must be unwound during the exoribonuclease halting event. We therefore hypothesized that the dianthovirus xrRNA structure observed in the crystal (with the P1 helix formed) is an important folding intermediate, but the RNA must change conformation in a programmed way to block Xrn1.

### An RNA pseudoknot interaction is necessary for Xrn1 resistance

The crystal structure suggests what the aforementioned putative conformational change could entail: Within the crystal lattice, nucleotides C43, U44, and C45 at the single-stranded 3′ end of one RNA molecule form Watson-Crick base pairs with G20, A19 and G18 in the loop region (L2A) of an adjacent RNA molecule (Fig. S4*A*). The formation of these pairs *in trans* in the crystal suggests that they could form *in cis*, creating a pseudoknot (PK) between the L2A and S2 regions (Fig. 2*D*). The potential to form this pseudoknot is present in all three dianthovirus xrRNA sequences (Fig. S4*B*), and in fact this putative pseudoknot can be extended by one additional Watson-Crick base pair (C17-G46; not present in the crystallized RNA). Indeed, addition of G46 to the 3′ end of the SCNMV xrRNA increased Xrn1 resistance *in vitro* and prompted us to perform all subsequent experiments using this 44 nucleotide-long xrRNA sequence (Fig. S4*C*). To determine the importance of the L2A-S2 pseudoknot, we mutated either L2A or S2 in SCNMV, RCNMV and CRSV xrRNA and assessed Xrn1 resistance *in vitro* (Fig. 2*E*, Fig. S4*D*). In all, disruption of the putative pseudoknot led to loss of Xrn1-resistance, whereas compensatory mutations (PK_COMP_) partially rescued Xrn1 resistance. We tested the importance of the pseuodknot for the accumulation of dianthovirus SR1f RNA by infecting *Nicotiana benthamiana* with RNA transcripts generated from an RCNMV RNA-1 infectious clone with either the WT, PK_S2_ or PK_COMP_ xrRNA sequences. Matching the *in vitro* data, disrupting the PK caused a complete loss of SR1f RNA formation, whereas compensatory mutations partially rescued production (Fig. 2*F*). We also determined that the PK interaction does not function *in trans* as dimer (Fig. S5*A*, *B*). These results show that the dianthovirus xrRNAs require the L2A-S2 pseudoknot interaction to block Xrn1.

### xrRNA pseudoknot formation creates a protective loop around the 5′ end

To understand how L2A-S2 pseudoknot formation could lead to exoribonuclease resistance, we constructed a model with this interaction formed *in cis*, and the P1 helix unwound, mimicking the structure at the moment of Xrn1 halting (Fig. 3*A, B*). In this modeled conformation, the 3′ end of the dianthovirus xrRNA encircles the 5′ end, thus creating a topology that is similar to the protective ring-like fold of mosquito-borne flaviviral xrRNAs (Fig. 3*B, C*). Thus, these xrRNAs appear to rely on a similar overarching strategy to block Xrn1, which is remarkable and unexpected given their very different sequences. In both cases, an apical loop forms long-range base-pairs with a downstream element, causing intervening sequence to loop protectively around the 5′ end (Fig. S5*C*). In the flavivirus xrRNAs, the loop is formed by two intact helical elements, while in the dianthovirus xrRNAs the P1 helix must be unwound. Comparison of these two xrRNAs shows the ring-like structure to be an emerging theme of RNA structure-based 5′→3′ exoribonuclease resistance, and convergent evolution can select different RNA sequences and structures to achieve this topology.

**Fig. 3.**
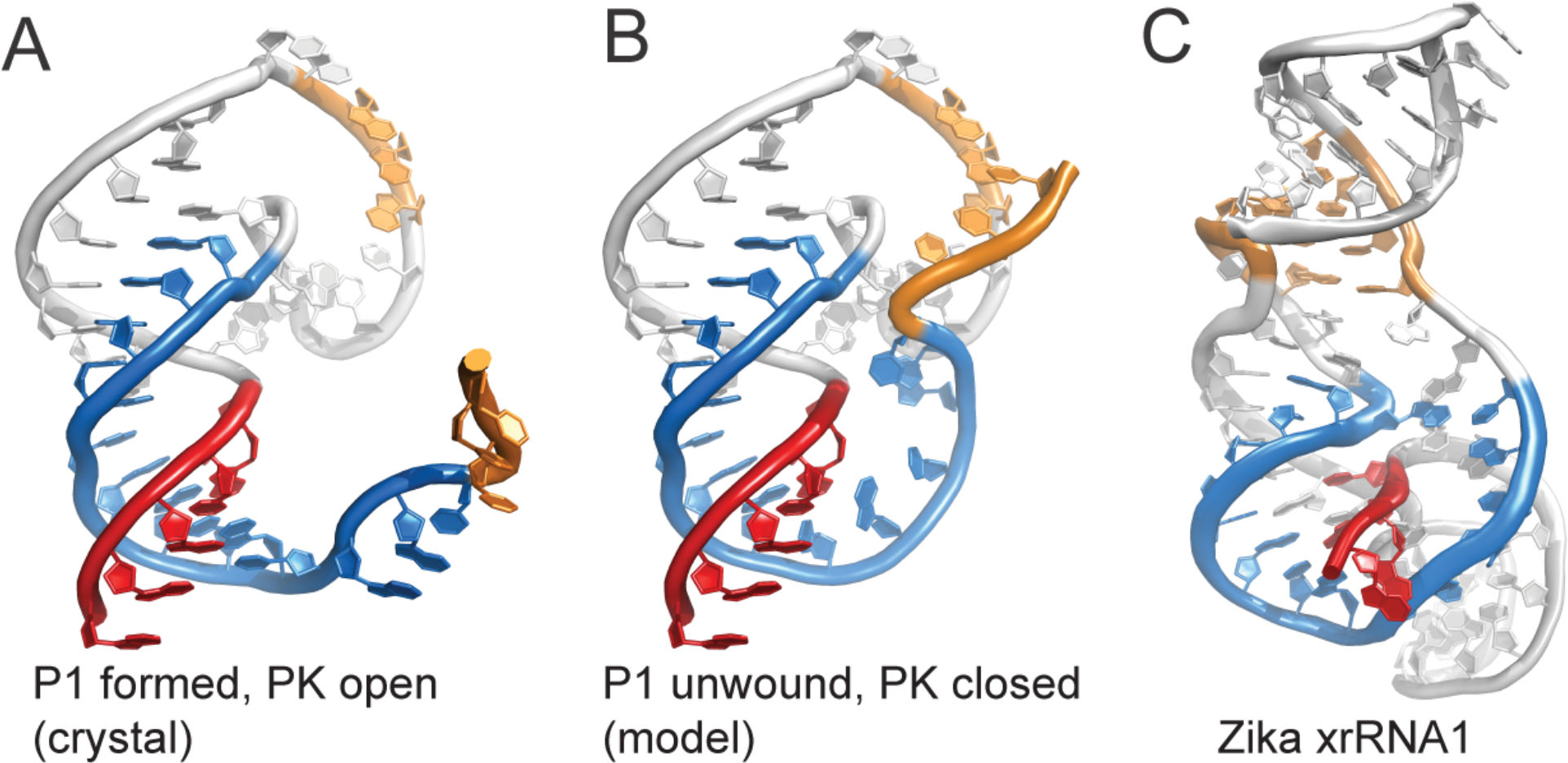
Pseudoknot formation creates a protective ring around the 5′end. (*A*) Crystallized SCNMV xrRNA. (*B*) Modeled xrRNA pseudoknot (PK) conformation (P1 unwound, PK formed). PK in orange. The 3′ end (blue) encircles the 5′ end (red). (*C*) Structure of ZIKV xrRNA1 (PDB: 5tpy) with 5′ end (red), ring-like fold (blue) and PK (orange) matching colors in *A-B*.

The SL conformation with P1 paired likely represents a folding intermediate to position S2 in such a way that when the PK forms it will encircle the 5′ end of the RNA. This interpretation suggests that the SL-to-PK switch itself is promoted by the tilted SL conformation created by tertiary interactions formed by the highly conserved sequence (Fig. S6*A*). One such interaction is enabled by L2B’s conformation, which is essentially a smaller loop embedded in the larger L2 loop. The backbone of L2B is similar to the U-turn found in some RNA tetraloops, but the loop is closed by a reverse Watson-Crick base pair between U21 and A28, with the latter base extruded from the P2 helix (Fig. S6*B*). The L2B structure creates a ‘docking point’ for A7, which is extruded from the L1 internal loop to stack between A25 and A26, thus inducing the overall tilted conformation (Fig. S6*C*). A similar long-range hairpin-adenosine interaction is found in 16S and 18S rRNA (Fig. S7). The importance of this interaction and thus the overall tilted SL conformation is revealed by an A7U mutant, which abolishes Xrn1 resistance *in vitro* and prevents SR1f RNA production uring RCNMV infection (Fig. S6*D*, *E*).

### Conformational dynamics of dianthovirus xrRNA

Our model suggests that formation of the essential S2-L2A pseudoknot involves a conformational switch between the SL and PK states. To test this we employed singlemolecule Förster resonance energy transfer (smFRET). Using the crystal structure and the PK model as a guide, we designed an RNA with dye modifications at sites that would result in high FRET if the PK was formed and a mid-FRET state if not (Fig. 4*A*, Fig. S8*A*). We verified that SCNMV xrRNA labeled with acceptor dye (Cy5) at position U15 was Xrn1-resistant (Fig. S8*D*), then annealed this RNA to a surface-immobilized DNA oligo labeled with a donor dye (Cy3) and monitored FRET efficiency by total internal reflection fluorescence (TIRF) microscopy. Wild-type SCNMV xrRNA predominantly exists in two states with ~0.8 and ~0.4 FRET efficiency (Fig. 4*B, C*). To assign these high-and mid-FRET states to specific conformations, we used the PKS2 mutant in smFRET experiments; it predominantly adopts the mid-FRET-associated conformation (Fig. 4*D, E*). Interestingly, smFRET traces following individual molecules over time demonstrate two types of wild-type SCNMV xrRNAs: RNAs ‘locked in’ a high FRET state and RNAs that undergo rapid conformational dynamics between the two states (Fig. 4*C*, Fig. S8*B*). This suggests two configurations in the PK state, indistinguishable by FRET: an unstable PK, which rapidly transitions back into the SL state, and a stable PK which remains folded for a longer period (perhaps stabilized by additional interactions). A subset of PK_S2_ molecules display a very rapid exchange between mid- and high-FRET states (Fig. 4*E*, Fig. S8*C*), but none remain in the high-FRET state; these molecules may sample the PK conformation but revert back to the SL state.

**Fig. 4.**
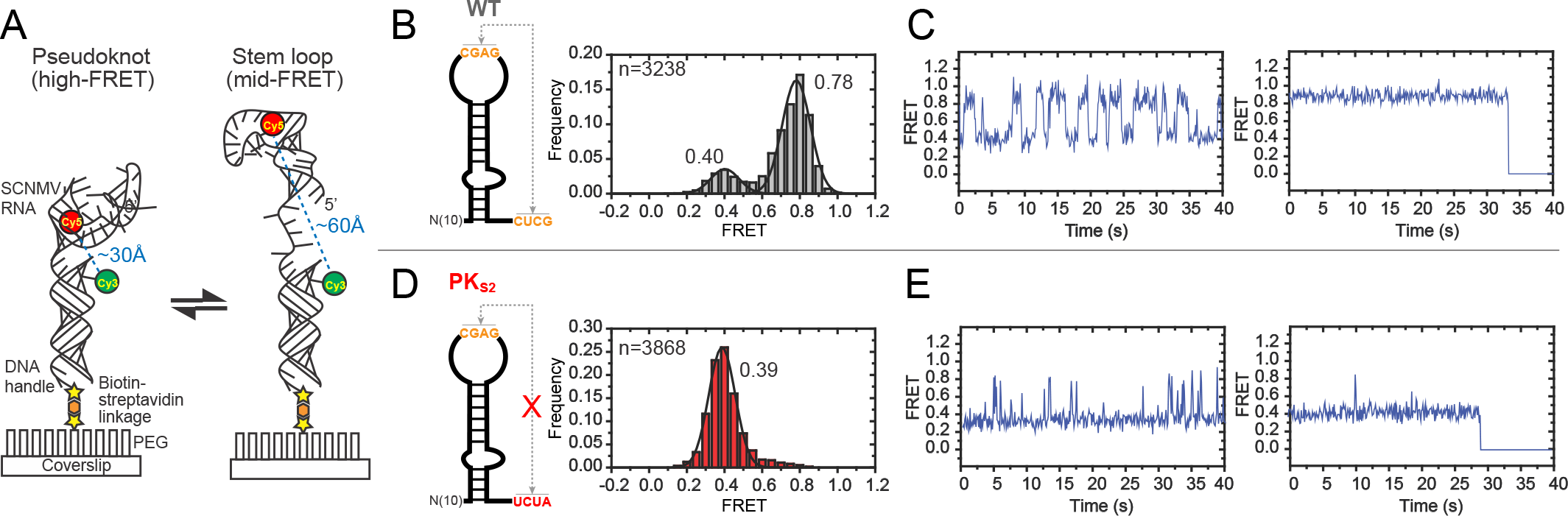
Conformational dynamics of SCNMV xrRNA. (*A*) Scheme of smFRET labeling strategy. (*B*) smFRET histograms of WT xrRNA. smFRET construct on the left. Average FRET values are shown next to peak. n = number of molecules represented by histogram. (*C*) Representative smFRET traces of individual WT xrRNA shows two separate populations. Left: dynamic population. Right: stable high-FRET population. (*D*) smFRET histograms of PK_S2_ xrRNA. Scheme of smFRET construct on the left. (*E*) Representative smFRET traces of two individual PK_S2_ xrRNA molecules.

### Co-degradational remodeling of dianthovirus xrRNA contributes to Xrn1 resistance

Dianthovirus’ Xrn1 resistance relies on the SL to PK conformational change, yet only the PK state can block Xrn1. These observations raise questions: What happens if Xrn1 encounters an xrRNA in the SL conformation? Is it able to quickly degrade the RNA or does the xrRNA refold in the presence of the exoribonuclease? Degradation-coupled unwinding of the P1 stem could potentially liberate the 3′ end of the RNA structure, favoring the formation of the PK and inducing Xrn1 resistance through ‘co-degradational’ RNA remodeling. If true, this model requires that RNA remodeling be faster than Xrn1-induced RNA hydrolysis. To test how an RNA in the SL conformation responds to Xrn1 degradation, we constructed an xrRNA with a 7-nucleotide 5′ extension that captures the 3′ end of the RNA in an extended P1 stem (P1EXT), preventing PK formation (Fig. 5*A*, Fig. S8*E*). As expected, the P1EXT xrRNA almost exclusively adopts a mid-FRET state corresponding to the SL conformation, indicating that the PK is not formed in this mutant (Fig. 5*B*). However, P1EXT is Xrn1-resistant (Fig. 5*C*), suggesting that the SL-to-PK switch required for exoribonuclease resistance is triggered during degradation, likely through unwinding of P1. Consistent with this, treating P1_EXT_ xrRNA with Xrn1 and examining the resultant RNA product by smFRET shows a redistribution of molecules from mid-FRET to high-FRET state (Fig. 5*D*). This demonstrates that the enzyme has shifted the xrRNA’s conformation from the SL to the PK state. Treatment with a catalytically inactive Xrn1 did not result in this redistribution (Fig. 5*E*), thus the enzyme’s degradation-linked helicase activity is necessary for the observed conformational change. Specifically, this result strongly suggests that unwinding of the P1 stem liberates the 3′ end of the structure to encircle the 5′ end and form the PK. These data show that Xrn1 can be co-opted into a ‘pseudo-chaperone’ function to favor the structure that ultimately prevents enzyme progression (Fig. 5*F*)

**Fig. 5.**
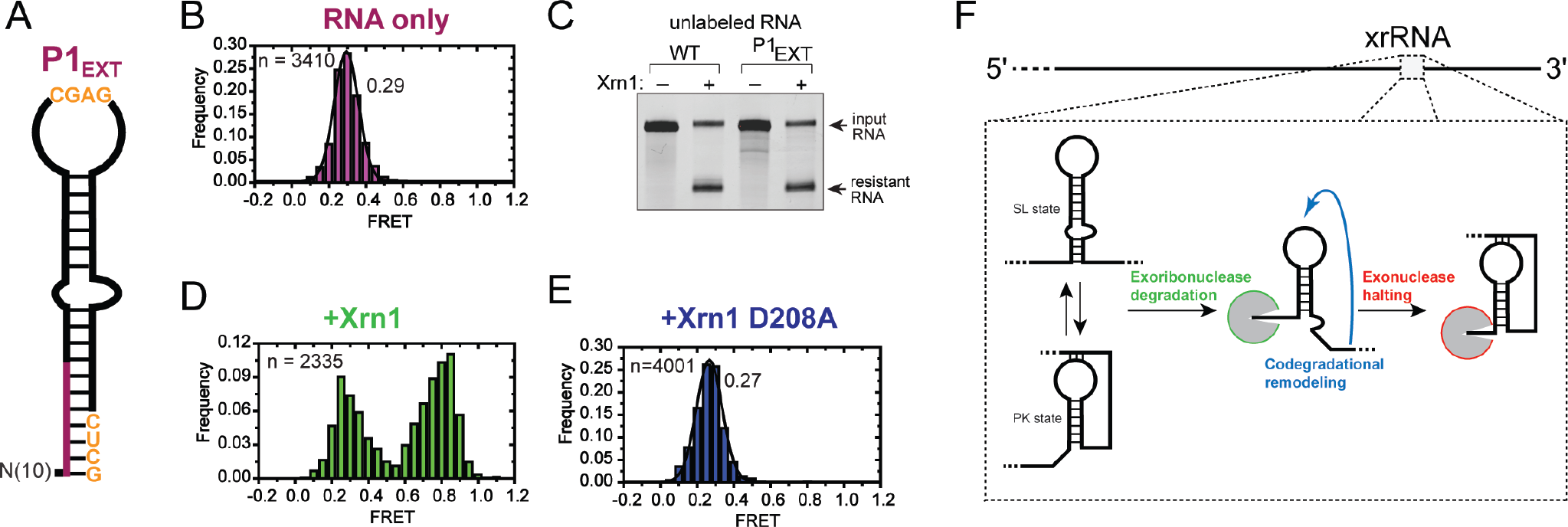
Co-degradational remodeling of SCNMV xrRNA structure. (*A*) Schematic representation of xrRNA smFRET construct with extended P1 stem (P1EXT). (*B*) smFRET histograms of P1EXT xrRNA. Average FRET value is shown next to peak. n = number of molecules represented by histogram. (*C*) *In vitro* Xrn1 degradation assay of WT and P1EXT xrRNA. (*D*-*E*) smFRET histograms of P1EXT xrRNA pretreated with WT Xrn1 (D) or mutant Xrn1(D208A) (*E*). n = number of molecules represented by histogram. (*F*) Scheme of the co-degradational remodeling of SCNMV xrRNA.

## Discussion

Exoribonuclease resistance as a means to generate non-coding RNA from larger precursors is an emerging theme in virology (21-24, 26). We have found that exoribonuclease-blocking sequences from plant-infecting dianthoviruses are authentic xrRNAs, generating viral non-coding RNAs in a process dependent on a specific but dynamic three-dimensional RNA fold. Furthermore, these xrRNAs appear to use the RNA unwinding activity of the exoribonuclease to stabilize their active structure in a form of ‘co-degradational remodeling.’ These findings establish xrRNAs as a distinct functional class of RNAs, and suggest RNA structure-dependent exoribonuclease resistance is a general mechanism of RNA maturation.

Flaviviruses and dianthoviruses belong to unrelated virus families, do not infect the same host species, and the xrRNA sequences have no similarity. Nonetheless, in both a folded xrRNA in the 3′UTR of a longer viral RNA is used to produce shorter non-coding RNAs that then play multiple (but different) important roles during viral infection (8–17, 26). Thus, a similar RNA maturation strategy has emerged in very divergent viruses, attesting to its utility and potential pervasiveness. Xrn1 can degrade most cellular RNAs, even highly structured ones (2); the fact that discrete xrRNA elements can stop the enzyme suggests that they have characteristics distinct from most folded RNAs. Indeed, while the three-dimensional structures of class 1 flaviviral xrRNAs and dianthoviral xrRNAs are different, they have similarities that suggest a shared overarching strategy. Both flaviviral and dianthoviral xrRNAs rely on a pseudoknot, a common RNA fold with many different configurations (28–30). However, in these xrRNAs, the pseudoknot creates a ring-like fold around the 5′ end of the structure, a topology not observed in other pseudoknot structures. The ring is comprised of sequences within the pseudoknot and downstream of the protected 5′ end, hence exoribonuclease hydrolysis-coupled helicase activity is unable to access or unfold the structure when the ring braces against the enzyme’s surface (7). Additionally, this shared topology suggests a way to confer directionality; the protective ring poses an obstacle to 5′→3′ decay, yet can be easily unwound by enzymes approaching from the 3′ side (such as the viral RNA-dependent RNA polymerase). Thus far, these ring-like folds have not been found in other RNAs and thus they might be a defining and necessary feature of xrRNAs.

Although flaviviral and dianthoviral xrRNAs share several similarities, there are important differences. In the flavivirus version, the fold requires a three-way junction, which interacts with the 5′ end directly to form a second pseudoknot interwoven with the first pseudoknot (7, 18). The stretch of RNA that forms the ring is almost entirely base-paired and a number of other tertiary interactions such as base triples and long-range non-canonical base-pairs stabilize the final fold (Fig. S5*C*). The dianthoviral xrRNAs are substantially shorter in length, there is no three-way junction, and most of the RNA that forms the ring appears to be single-stranded as a result of the unwinding of stem P1 (Fig. S5*C*). These differences reveal how very divergent sequences and secondary structures can both give rise to the ring-like topology. Because the crystal did not contain the dianthoviral xrRNA in the PK form, we do not yet know the full repertoire of tertiary interactions that stabilize this conformation; local remodeling of the structure could give rise to additional interactions. A specifically configured pseudoknot may be a common feature of all xrRNAs, but more structures are needed to test this and to understand other shared features.

The unusual nature of the xrRNA fold invites speculation in regard to how it forms correctly; RNA pseudoknots are ubiquitous but only xrRNAs have been observed to form a protective ring. This implies that the high levels of Xrn1 resistance observed *in vitro* requires a mechanism to ensure that the ring always wraps around the 5′ end of the structure. In flaviviral xrRNAs, the structure shows that the 5′ end must interact directly with the three-way junction for the junction to fold and form the ring structure (19). In dianthoviral xrRNAs, the data suggest that the P1 helix initially forms to position the 5′ and 3′ ends correctly (SL state), then a programmed conformational change leads to P1 helix unwinding and pseudoknot formation (PK state). This pathway could help ensure that pseudoknot formation establishes the ring around the 5′ end. Accordingly, mutations affecting the putative folding intermediate (i.e. mutations to the P1 stem) leads to a ~50% reduction in Xrn1 resistance; the most straightforward interpretation is that failure to form the P1 stem decouples pseudoknot and ring formation from 5′ end placement and ~50% of the molecules are misfolded.

Additional evidence for the folding model described above is that dianthoviral xrRNAs retain their activity even when the PK state is prevented from forming through an extended P1 stem. This observation and the shift toward the PK state when Xrn1 degrades the RNA reveals that RNA remodeling can happen ‘co-degradationally’ and that the conformational switch between the SL and PK states is faster than the progression of the enzyme through the structure. The helicase activity of Xrn1 can thus be co-opted into a pseudo-chaperone function: by unfolding P1, Xrn1 liberates nucleotides required for pseudoknot formation and shifts the conformation to the resistant state. Although smFRET shows that pseudoknot formation can occur spontaneously *in vitro*, within the cell and in the context of the full-length viral RNA, co-degradational remodeling may be an important. It could allow the xrRNA to respond to the RNA decay machinery while staying flexible enough for 3′→5′ unfolding by the viral RNA-dependent RNA polymerase. To our knowledge, there are no other known examples of co-degradational RNA structure formation or remodeling, but our discovery suggests that similar pseudo-chaperone function dependent on exoribonucleases or helicases may exist elsewhere.

Exoribonuclease resistance conferred by RNA structure may be a more common mechanism than hitherto anticipated, and recent studies have identified putative xrRNAs in several virus families. A conserved sequence that blocks 5′→3′ exoribonucleolytic decay was recently found in the 3′UTR of plant-infecting benyviruses and cucumoviruses; both have multipartite positive-sense RNA genomes (23, 24, 31). Moreover, Xrn1 is confounded by RNA elements in the 5′UTRs of hepaci- and pestiviruses (21), as well as by a G-rich sequence structure in the N mRNA of Rift Valley fever virus, and multiple RNA structures in ambisense-derived transcripts of arenaviruses (22). There are no obvious common sequence patterns and most await detailed functional and structural characterization.

The range of biological functions of ncRNAs produced by xrRNAs (8–17, 23, 26), as well as their presence in diverse virus families, suggests that regulated exoribonuclease resistance could be a pathway for RNA maturation in cells. Of note, G-rich RNA sequences installed artificially into mRNAs have been shown to confound Xrn1 activity in yeast (presumably through G-quadruplex structures) but the same G-rich sequences appear unable to stall exoribonucleolytic decay in mammalian cells (32). Moreover, pentatricopeptide repeat (PPR) proteins function as protein barriers for exoribonucleolytic decay and thus define mRNA termini in chloroplasts and mitochondria (33–35). Likewise, rRNA processing involves precise trimming by exoribonucleases (36, 37). It is conceivable that cellular xrRNAs might exist to generate biologically active 5′-truncated decay intermediates. Indeed, many RNA-centered processes used by cells were first discovered in viruses, including RNA structures that block 3′→5′ decay (38, 39). This illustrates how RNA elements identified in viruses can inform the search for related molecular structures elsewhere in biology and motivate ongoing research to understand the enormous complexity and diversity of structured RNAs.

## Materials and Methods

### RNA transcription and protein expression

RNA was *in vitro* transcribed with T7 polymerase and purified by denaturing polyacrylamide gel electrophoresis. *Kleuveromyces lactis* Xrn1 (4) (residues 1-1245), *Kleuveromyces lactis* Dxo1 (40), *Bacillus subtilis* RNase J1 (41) and *Bdellovibrio bacteriovorus* RppH (42) were 6XHis-tagged, expressed in *E. coli* BL21 cells and purified using Ni-NTA resin (Thermo) and size exclusion chromatography. For details see *SI Materials and Methods*.

### Exoribonuclease degradation

Xrn1 resistance assays were as previously described (6). Briefly, RNA was refolded then incubated with recombinant RppH and recombinant exoribonuclease (Xrn1, Dxo1, RNase J1) for 2 hours at 30°C. Reactions were resolved on a denaturing polyacrylamide gel. For quantitative Xrn1 resistance assays, RNAs were radiolabeled at their 3’ terminus and Xrn1 resistance was defined as the percentage of Xrn1-resistant degradation products (resistant RNA) relative to the total amount of untreated RNA (input RNA). For details see *SI Materials and Methods*.

### Mapping of the Xrn1 stop site

Xrn1-resistant degradation products were reverse transcribedwith a FAM (6-flourescnetly labeled sequence-specific reverse primer and the cDNA analyzed by capillary electrophoresis. For details see *SI Materials and Methods*.

### Mutagenesis of RCNMV infectious clone and viral infection

Mutations were introduced into an infectious cDNA construct of RCNMV RNA-1 (27) using a PCR-based method. RNA-1 was transcribed in vitro and then combined with an equal volume of wild-type RCNMV RNA-2 transcripts plant inoculations. Carborundum-dusted leaves of *Nicotiana benthamiana* were rub inoculated with RCNMV transcripts and the infection allowed to proceed for 3 days. Total RNA was extracted from inoculated leaves. For details see *SI Materials and Methods*.

### Northern blotting

RNA from mock- or RCNMV-infected cells was resolved on denaturing PAGE, transferred to a HyBond-N+ nylon membrane, UV crosslinked and blocked. Blots were probed with a [5’-^32^P]-labeled 35-mer DNA, washed four times, then phosphorimaged. For details see *SI Materials and Methods*

### RNA crystallization and structure determination

RNA was crystallized in 50 mM sodium cacodylate trihydrate pH 6.5, 0.2 M potassium chloride, 0.1 M magnesium acetate tetrahydrate, 14% w/v PEG 8000. Crystals were buffer-exchanged into cryo-colling solution (above buffer with 20% ethylene glycol and 20 mM Iridium (III) hexamine) and flash-frozen in liquid nitrogen. Diffraction data were collected at Advanced Light Source Beamline 4.2.2 using ‘shutterless’ collection. Data were indexed, integrated, and scaled using XDS (43). Nine Iridium (III) hexammine sites were identified and used in SAD phasing. Model building was done in COOT (44, 45) and Phenix (46). Data was collected and processed to 2.55 Å, but only data to 2.9 Å were used for refinement. Detailed information in *SI Materials and Methods* and Table S1.

### Single-molecule FRET

Cy5-labeled purified RNAs were annealed to biotinylated Cy3-labeled DNA oligo and immobilized on PEGylated cover slides (Ted Pella, Inc.). Slides were imaged on a total internal reflection fluorescence microscope (Nikon Eclipse Ti-E). Analysis was performed using custom MATLAB (Release 2017a, The MathWorks) scripts. FRET is defined as the ratio of I_A_/(I_A_+I_D_), where I_A_ is the acceptor intensity and ID is the donor intensity. Histograms were compiled from the average FRET value obtained from molecules over a 2-sec observation time. Molecules with 0.0 FRET, obtained due to a fraction of molecules containing bleached acceptor fluorophores, were excluded from histograms by omitting molecules that displayed lower than a minimum threshold value of acceptor intensity. For details see *SI Materials and Methods*

## Acknowledgments

We thank C. Musselman and O. Rissland for critical reading of this manuscript. Support from NIH grants R35GM118070 and R01AI133348 (J.S.K.); EMBL Fellowship ALTF 611-2015 (A.-L.S.); NIH F32GM117730 (B.M.A.). The UC Denver X-ray Facility is supported by grants P30CA046934 and S10OD012033. The ALS is supported by the Director, Office of Science, Office of Basic Energy Sciences of the U.S. Department of Energy under Contract No. DE-AC02-05CH11231. T.L.S. acknowledges the support of the North Carolina Agricultural Research Service in the College of Agriculture and Life Sciences at North Carolina State University.

